# Convolutional models of RNA energetics

**DOI:** 10.1101/470740

**Authors:** Michelle J. Wu

**Affiliations:** Biomedical Informatics Training Program, Stanford University, Stanford, CA 94035

## Abstract

Nucleic acid molecular biology and synthetic biology are undergoing rapid advances with the emergence of designer riboswitches controlling living cells, CRISPR/Cas9-based genome editing, high-throughput RNA-based silencing, and reengineering of mRNA translation. Many of these efforts require the design of nucleic acid interactions, which relies on accurate models for DNA and RNA energetics. Existing models utilize nearest neighbor rules, which were parameterized through careful optical melting measurements. However, these relatively simple rules often fail to quantitatively account for the biophysical behavior of molecules even *in vitro*, let alone *in vivo*. This is due to the limited experimental throughput of optical melting experiments and the infinitely large space of possible motifs that can be formed. Here, we present a convolutional neural network architecture to model the energies of nucleic acid motifs, allowing for learning of representations of physical interactions that generalize to arbitrary unmeasured motifs. First, we used existing parameterizations of motif energies to train the model and demonstrate that our model is expressive enough to recapitulate the current model. Then, through training on optical melting datasets from the literature, we have shown that the model can accurately predict the thermodynamics of hairpins containing unmeasured motifs. This work demonstrates the utility of convolutional models for capturing the thermodynamic parameters that underlie nucleic acid interactions.

## 1 Introduction

RNA has recently emerged as the ideal candidate for a building block to create the next generation of therapeutics, with the potential to specifically disable or manipulate the genes involved in disease. RNA’s functional versatility is exemplified by the development of myriad novel RNA-based synthetic biology tools, including designer riboswitches controlling living cells [24, 31, 8], CRISPR/Cas9-based genome editing [13, 17], RNA-based silencing [20, 22], and reengineering of mRNA translation [21, 7]. The ability to design RNA elements is central to gaining precise control over these systems and customizing them for use as potential RNA-based therapeutics. However, doing so relies upon a thorough understanding of the energetics of RNA interactions, and computational models have proven inadequate to quantitatively account for the biophysical behavior of RNA molecules. This lack of predictive models has been a major barrier to the wide adoption of existing RNA-based technologies.

Current nucleic acid energetic models have relied upon nearest neighbor rules, which posit that the thermodynamic contribution of a base pair depends only upon the sequence of itself and its nearest base pair [2, 25, 34]. This enables the energy of a given structure to be estimated by breaking it up into its constituent motifs. Dynamic programming techniques can be applied to estimate the lowest energy conformations of a given sequence [23].

While this model has been sufficient as a guide for RNA secondary structure stabilities, it falls short in painting a complete picture of an RNA molecule’s full conformational ensemble. This is largely because the model has been parameterized using less than 10^3^ optical melting measurements [27, 35, 34, 18] and rough, sequence-agnostic approximations are used for unmeasured motifs. While optical melting experiments have been adequate to parameterize base pair energies, more careful modeling is necessary to characterize the loops in the intervening regions. These secondary structure elements are key to positioning RNA helices in the correct orientation and determining the overall structure.

A major challenge in modeling these loop elements is that the sequence space for loop elements, including hairpin loops, internal loops and multiloops, grows exponentially with length. Thus, accurate energetic models require parameterization of this space without the ability the make direct measurements of every element. State-of-the-art nearest neighbor models use linear approximations based on the length of the loop, with terms for the identity of the closing base pair and first mismatch for hairpin loops [12, 27]. However, this model clearly ignores important sequence-dependent energetic contributions that are exacerbated as loop sizes increase [33, 32].

Convolutional neural networks (CNNs) are a particularly suitable approach for extracting the sequence contributors to these energies from massive experimental datasets. These models are hierarchical in nature, mirroring the folding process of complex RNA structures [4]. In addition, they have been shown to be effective in other tasks involving modeling of nucleic acid sequences [1, 37, 14]. Thus, convolutional models may be an effective architecture for modeling the interactions between neighboring bases that will allow for generalization of the energetic model to large loop elements. Here, we explore the use of CNNs for predicting the energies of unmeasured RNA structural motifs.

## 2 Methods

### 2.1 Datasets

Initially, models were trained directly on motif energies given by parameter sets for the nearest neighbor model [18]. These were parsed from the files used by the NUPACK software [36]. The dataset includes Watson-Crick helices, triloops and tetraloops, mismatches, dangling ends, and 1×1, 1×2, and 2×2 internal loops. In addition, 1,000 multiloops were randomly generated and their energies were calculated based on the linear approximation described by Turner and colleagues [18]. This resulted in a set of 13,207 motifs, which were split into a training and test set.

Models were then trained and evaluated on experimental data from the literature. These data were taken from six different papers that made optical melting measurements for 109 RNA hairpins at standard conditions of 1M NaCl [9, 10, 28, 26, 27, 19]. 28 of the hairpins with the largest loop elements, with seven or more unpaired nucleotides, were held out as test data to evaluate the ability of the model to generalize to larger loops, and 10 of the remaining training hairpins were used as a validation dataset for hyperparameter tuning. An additional dataset of 33 hairpins taken after the training of the nearest neighbor parameter sets was used as an additional test set [30]. These papers all used standard optical melting fitting procedures, and Δ*G*_37_ values (folding free energy at 37°C) reported by each paper were used for training.

### 2.2 Model training

For the experimental data, since motif energies cannot be individually measured, the model must be trained from energies for an entire structure. To simplify training, each hairpin was assumed to only adopt a single secondary structure, as specified in the original papers, and undergo a two-state transition between folded and fully unfolded. These assumptions enable us to easily decompose each hairpin into its subcomponent motifs (Figure 1). As in the classical nearest neighbor model, the overall energy is modeled as the sum of individual motif energies, and the gradients were backpropagated through this sum during training.

**Figure 1:**
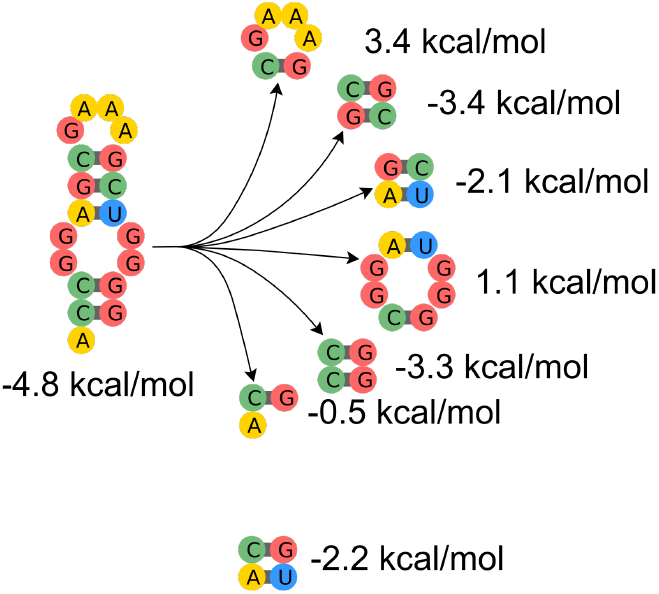
Nearest neighbor model for RNA energetics. Each secondary structure is broken up into overlapping structural motifs, whose folding free energies are assumed to be additive.

During each training step, a random batch of hairpins is selected and a single motif length is selected to simplify the tensor operations in training. Motifs in our dataset range from length 2 to 11, and a length is also chosen proportionally to the number of motifs of that length.

### 2.3 Model architecture

As in the nearest neighbor model for RNA energetics, each secondary structure is decomposed into individual motifs (Figure 1) [3, 27, 35]. The CNN is used to predict the energy of each of these motifs, and the model assumes that these energies are additive.

Each motif is represented using a circular matrix, where pairs of bases that are linked by phosphodiester bonds along the backbone of the RNA or by hydrogen bonds in a base pairing interaction are consecutive rows in the resulting matrix (Figure 2, left). This ensures that the close proximity of interacting bases is captured by the structure of the matrix representation.

**Figure 2:**
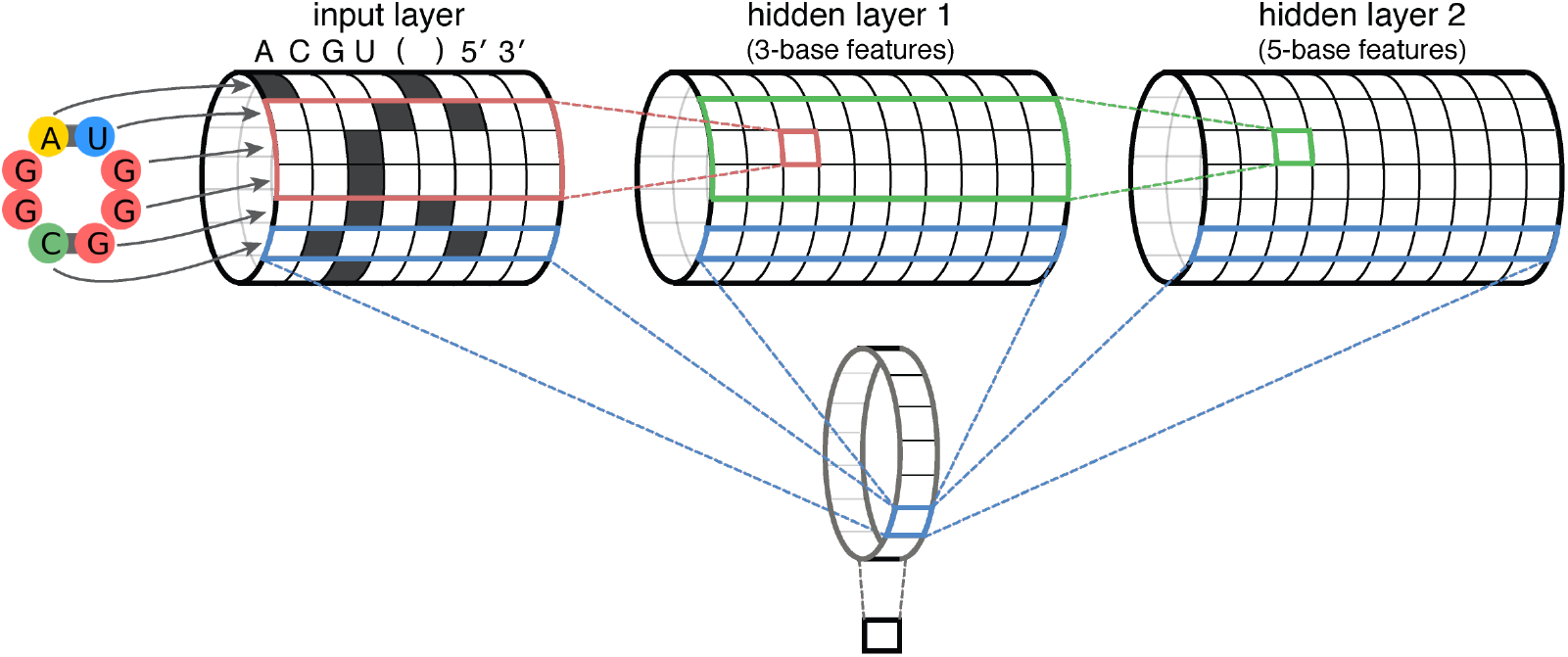
CNN model for RNA motif energies. Each motif is converted to a row within a binary matrix (left), with columns representing sequence identity, base pairing, and whether a base is a part of a 5′ or 3′ dangling end (as in the last motif in Figure 1). Convolutions are computed on this circular input matrix to get the first hidden layer (red) and similarly for subsequent hidden layers (green). These convolutional layers are able to represent increasingly long sequence features, with 3-base features in the first hidden layer, 5-base features in the second hidden layer, and so on. Finally, these features are aggregated into a final fully connected layer (blue) and summed to produce the output (gray).

Given this matrix representation of each motif, 1D convolutions with width 3 and stride 1 are applied over the sequence dimension (Figure 2, red & green). Multiple layers of convolutions can be applied, with the values in the first hidden layer representing sequences features involving up to 3 bases, the second layer up to 5 bases, and so on. This architecture allows the model to learn sequence features bottom-up, integrating shorter motifs into longer and longer motifs at each layer. Interactions between nucleotides that are physically near each other in space (e.g. base pairing) are accounted for in earlier layers, so that a nucleotide’s immediate context is taken into consideration as longer range interactions are represented in later layers. To get the final energy, the sequence features in all hidden layers are combined into a final fully connected layer (Figure 2, blue) and summed over all positions to get a single value regardless of length (Figure 2, gray).

For the motif-level model, we used 10 convolutional layers of 50 units each, while for the optical melting model, we used a single convolutional layer of 20 units. In both cases, we used a learning rate of 10^−4^ and a dropout rate of 0.5.

### 2.4 Software

Code for training and testing models is available at https://github.com/wuami/nn2.

## 3 Results

### 3.1 Recovering nearest neighbor energies with a CNN

To evaluate the expressivity of this model, we first applied the model to the individual motif energies given by the Turner 1999 parameter set [18]. This dataset encompassed all explicity parameterized values, including base pairs, apical loops, and internal loops. In addition, 1,000 multiloops were randomly generated and their energies were calculated based on the linear approximation described by Turner and colleages [18]. In total, this dataset comprised of over 13,000 motif energies.

We were able to train a model that captured this parameter set with an RMSE of 0.11 kcal/mol on a held out test set (Figure 3). Excluding the multiloops that are not explicitly parameterized, with energies derived directly from a linear model, the RMSE is 0.21 kcal/mol, comparable to the error of the parameter estimates given by the fit in the original paper [18]. The error was similar for motifs of different lengths and types. This suggests that the CNN model is at least as expressive as the original Turner mode and should be able to capture RNA energetic parameters at least as well. Furthermore, this result suggests that this model can be trained to capture generalizable motif features, allowing it to generalize to an unseen test set derived directly from experimental data.

**Figure 3:**
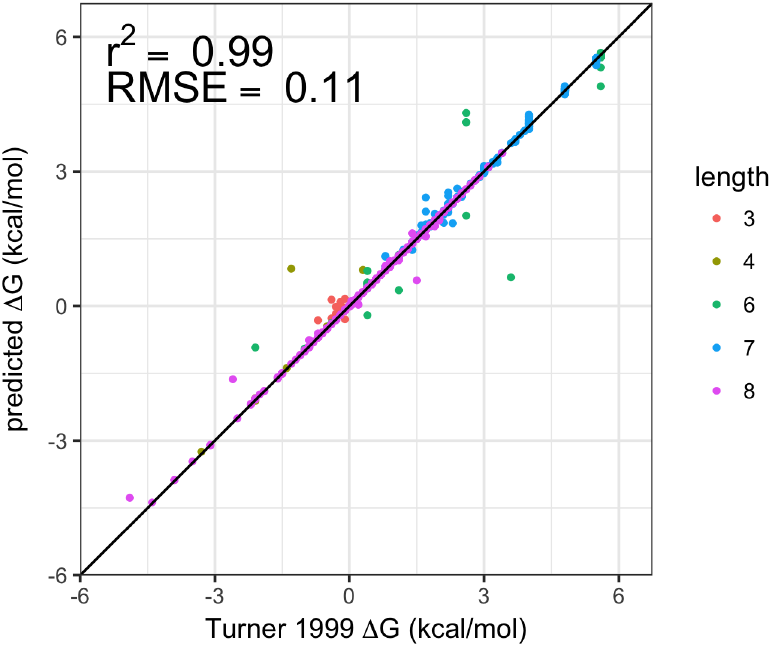
A CNN model accurately recovers the Turner 1999 [18] parameters on a test set for motifs of various lengths, including base pairs, internal loops, and multiloops.

### 3.2 Training models on optical melting data

To validate the effectiveness of this model trained directly on experimental data, we turned to optical melting data from the literature. These experiments are considered to be the gold standard in measuring nucleic acid thermodynamics [34, 25, 18]. Our dataset consisted of measurements for 109 RNA hairpins containing 125 unique motifs. Hairpins containing a loop of more than 6 nucleotides were held out as a test set to evaluate the model’s ability to generalize learned features to larger unmeasured motifs. A separate validation set from the shorter loop hairpins was used for hyperparameter tuning.

As a comparison, we made predictions for the same constructs using ViennaRNA, a commonly-used secondary structure prediction software package [16]. This package uses nearest neighbor parameters derived from a set of measurements that includes our data, effectively using our test hairpins in training of their model. Nevertheless, our model performs similarly to ViennaRNA as measured by correlation. In addition, we outperform ViennaRNA when excluding one paper for which neither model is able to make meaningful predictions. However, the CNN predictions are shifted by a constant offset, resulting in poor RMSE values (Figure 4). While this offset could likely be mitigated by training on a more comprehensive dataset, it should have no effect on predictions made with downstream minimum free energy secondary structure or partition function calculations, as long as all hairpins have the same error.

**Figure 4:**
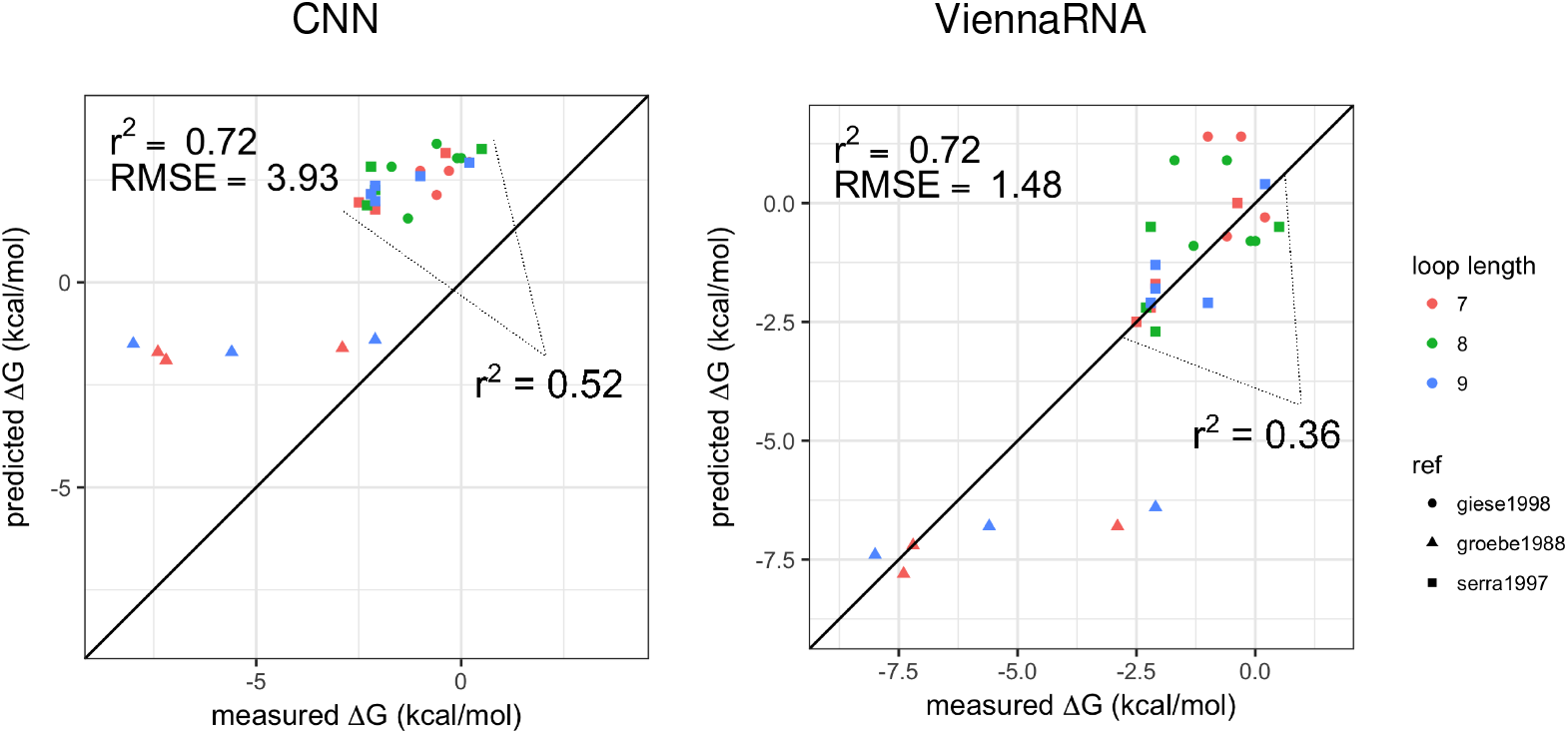
The CNN performs similarly to ViennaRNA in terms of correlation, despite the ViennaRNA model being trained partially on the test dataset.

In addition, we tested our model on a separate dataset of hairpins collected after the Turner 1999 parameters were published [30]. While our model performed worse than ViennaRNA overall, it is able to capture variance within hairpins with the same closing base pair (Figure 5). This suggests that, in combination with information from the existing nearest neighbor models, it may be able to capture energetic differences in loops across sequence variants with improved accuracy.

**Figure 5:**
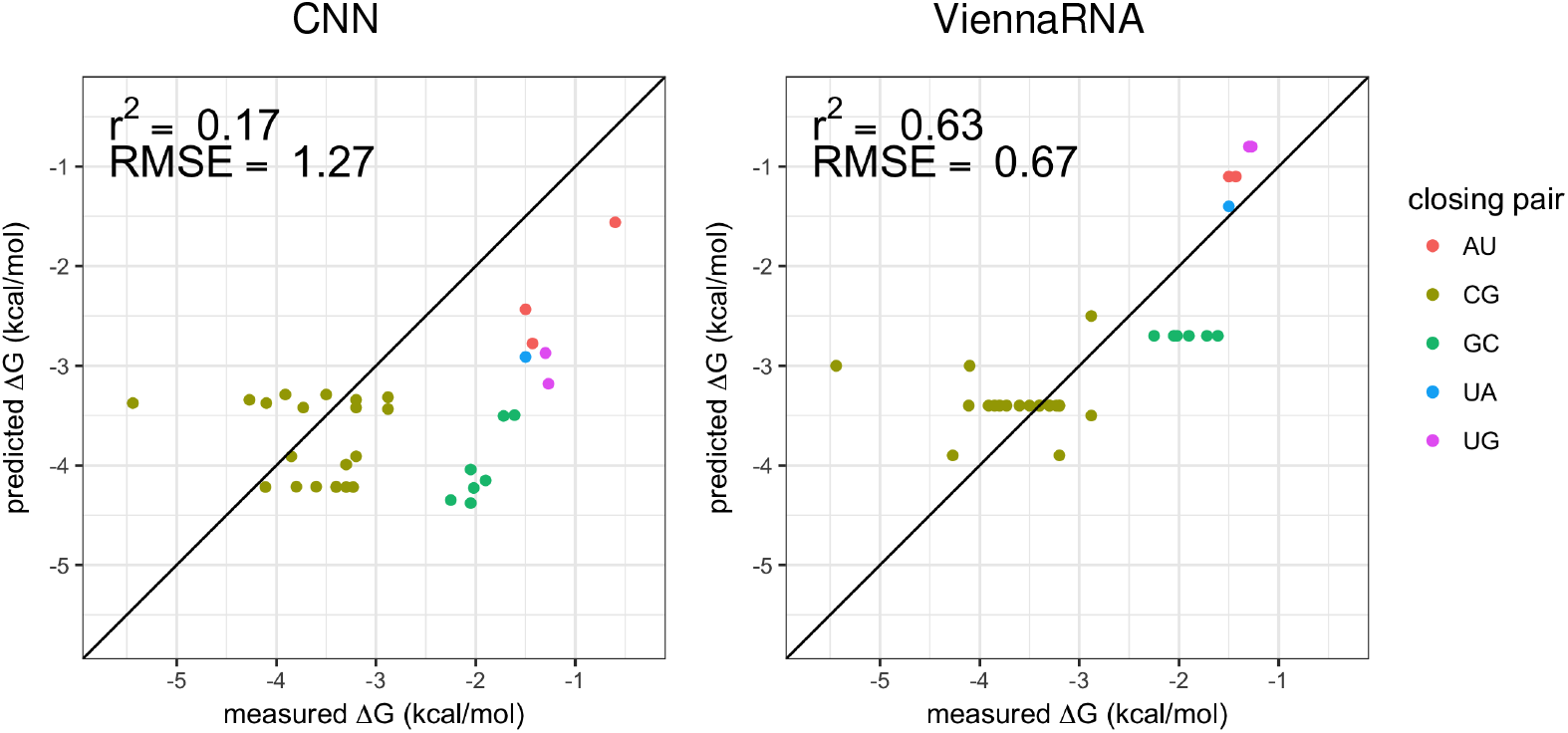
On a more recent dataset, the CNN performs worse overall but is able to capture variance within sets of hairpins with the same closing base pair.

The simple architecture of our model also enables some interpretation of the convolutional filters, which encode motif features that contribute to energetic predictions. Figure 6 shows the weights trained for the first convolutional layer, which can be interpreted as features that incorporate information from 3-mers of the motif. For example, the highest weighted convolutional filter (left) has strongly positive weights for a G and C that are adjacent in the motif representation, suggesting it may be a detector of GC base pairs, which would be expected to be a strong contributor to motif energies.

**Figure 6:**
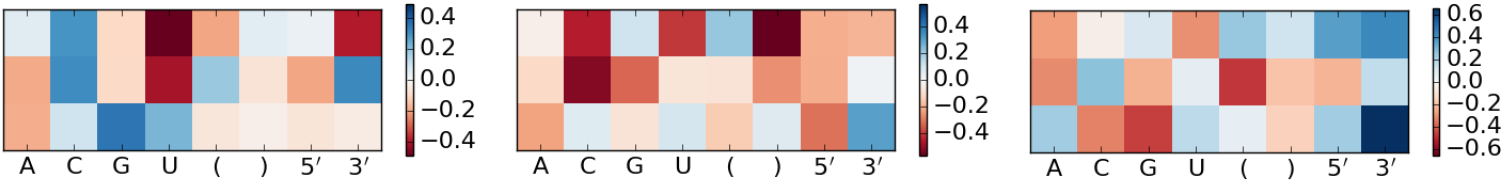
The three convolutional filters with the strongest weights in the fully connected layer show the input features contribute to predicting motif energies.

## 4 Discussion

Modern technologies have enabled massively parallel, quantitative measurement of tens of thousands of individual RNA sequences [5, 29, 15]. These methods will allow for high-throughput measurement of melting curves at the scale needed for more exhaustive characterization of motifs. However, given the arbitrarily large number of possible motifs that can be formed by nucleic acids, models that generalize learned principles to unmeasured motifs are still necessary. These rich datasets will enable the training of deeper and more expressive convolutional models. Due to the small size of our dataset, we only used a one-layer network for the experimental validation, limiting the ability of the model to represent higher order features. In addition, the constant offset observed in our predictions for hairpins containing long motifs would likely be resolved by a larger dataset containing a more diverse set of motif lengths.

Existing RNA secondary structure prediction software use dynamic programming methods to determine the lowest energy conformation that a given sequence can adopt. As a result of removing the linear assumptions used for multiloops in these algorithms, the dynamic programming algorithm also becomes infeasible. Integrating the CNN model into these methods would require computing the energies of all motifs that could be formed by a particular RNA. While many short motifs could be precomputed for the purposes of secondary structure prediction software, this may be infeasible for long RNA molecules, and computational shortcuts may be necessary to reduce the search space [11, 6].

This work demonstrates the utility of convolutional neural networks for modeling nucleic acid thermodynamics. Our models enable more accurate modeling of unmeasured motifs and illustrates the need for modern, high-throughput measurements to advance predictive models of nucleic acid thermodynamics.

## Acknowledgments

This work was supported by the National Science Foundation Graduate Research Fellowship Program under Grant No. DGE-114747 and an NIH Ruth L. Kirschstein National Research Service Award from the NIGMS (F31GM125151). This work used the XStream computational resource, supported by the National Science Foundation Major Research Instrumentation program (ACI-1429830).

